# Enzymatic attachment of antigens to probe immune modulation by skin commensals

**DOI:** 10.64898/2026.10.31.763323

**Authors:** Djenet Bousbaine, Pranav Lalgudi, Katherine D. Bauman, Eunice Tsang, Tam T. D. Nguyen, Anchaleena James, Miki Chang, Aishan Zhao, Masahiko Terasaki, Joyce M. Swenson, Anne Trinh, Michael A. Fischbach

**Affiliations:** Department of Bioengineering, Stanford University, Stanford, CA 94305, USA; ChEM-H Institute, Stanford University, Stanford, CA 94305, USA; Department of Genetics, Stanford University School of Medicine, Stanford, CA 94305, USA; Microbiome Therapies Initiative, Stanford University, Stanford, CA 94305, USA; Chan Zuckerberg Biohub, Stanford, CA 94305, USA

## Abstract

The skin colonist *Staphylococcus epidermidis* induces potent T and B cell responses, which can be redirected against antigens of interest by genetic engineering. However, since many taxa in the microbiome are genetically intractable, the utility of this method is limited. Here, we use recombinant sortase to conjugate antigens to the surface of a wide range of Gram-positive bacterial species, and we show that this non-genetic method can be used to generate protective immunity in vivo.

## MAIN TEXT

The skin colonist *Staphylococcus epidermidis* (*S. epi*) induces antigen-specific T cell responses upon colonization^1^. Building on this discovery, we demonstrated that colonization with *S. epi* led to the induction of a potent and pre-emptive antibody response^2,3^. Strains of *S. epi* engineered to display non-native antigens elicit therapeutic T and B cell responses against targets of interest^2,4^.

However, the methods described in these studies are only possible in organisms with genetic tools, rendering most members of the microbiome inaccessible. To solve this problem, we sought to develop an enzymatic strategy to conjugate antigens to the bacterial cell surface without the need for genetic tools (Fig. 1a). This approach was inspired by our observation that a strain of *S. epi* expressing SpyCatcher^5^ on its surface (Aap-*sc*), when conjugated to the model antigen tetanus toxin fragment C (Aap-*sc*-TTFC), elicited equivalent antibody responses to a strain expressing the same antigen constitutively (wTTFC)^2^. Despite the transient nature of the antigen exposure, colonization with either wTTFC or Aap-*sc*-TTFC led to long-lived antibody responses both systemically and at mucosal sites, months after the last colonization event (Extended Data Fig. 1). This suggests that short-term display of antigens on microbes is sufficient to induce potent and long-lived antibody responses in vivo.

**Figure 1:**
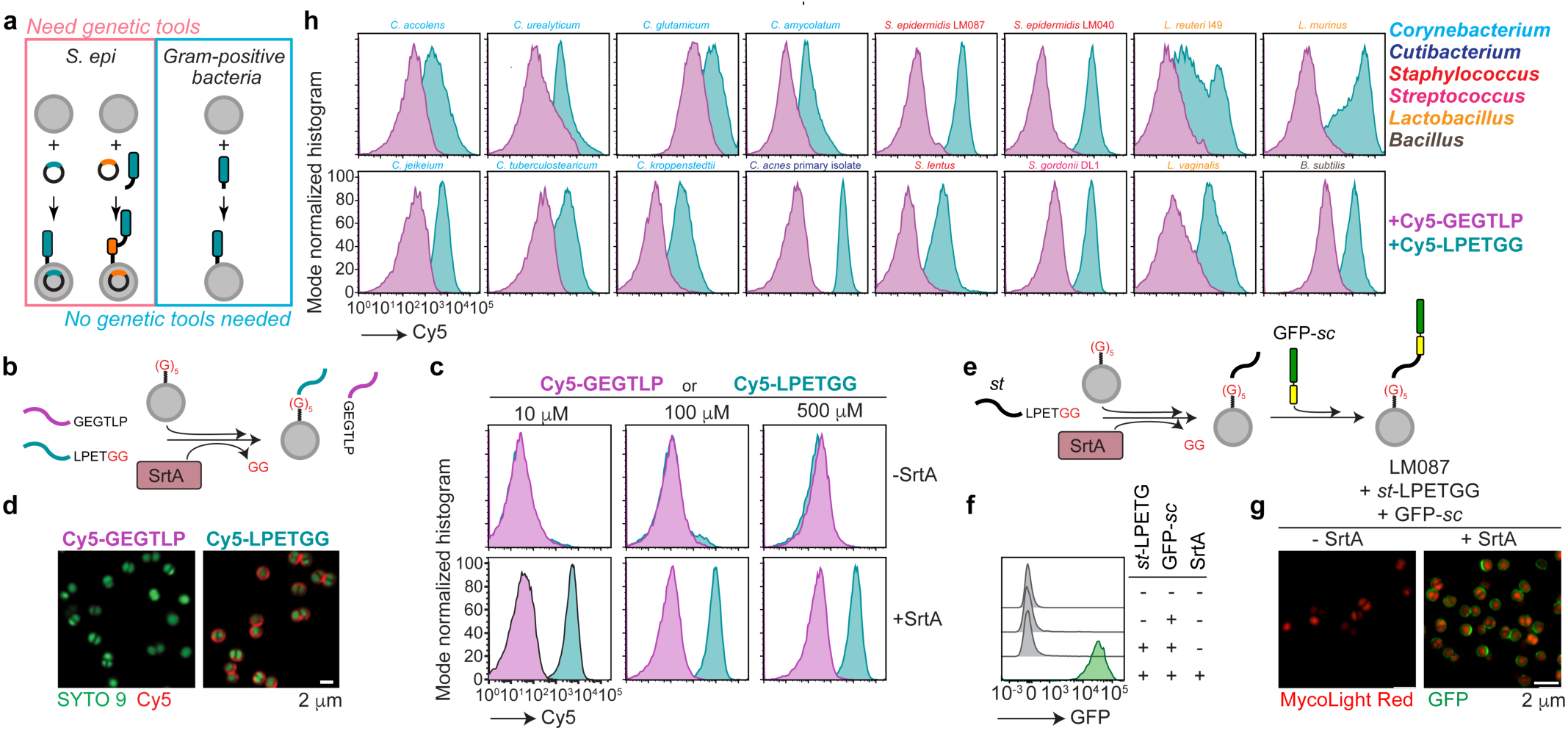
Sortase-mediated labeling of gram-positive bacteria with peptides and proteins. (**a**) We previously developed tools to display antigens on the surface of *S. epidermidis* (*S. epi*) via genetic engineering for topical vaccination (pink). Here, we developed a sortase A (SrtA)-mediated method to display peptide and protein antigens to the surface of Gram-positive bacteria, without the need for genetic tools (blue). (**b**) Schematic of SrtA-mediated labeling of bacteria with peptides. Cy5-containing peptides are equipped with a SrtA recognition motif (LPETGG) or a scramble motif (GEGTLP) and mixed with bacteria and recombinant SrtA. (**c**) *S. epi* LM087 was incubated with varying amounts of Cy5-containing peptides in the presence or absence of 50 μM SrtA. Peptide conjugation was monitored by flow cytometry. (**d**) Representative microscopy images of SrtA labeling of *S. epi* LM087 with 500 μM of the indicated peptide and 50 μM of SrtA. (**e**) Schematic showing SrtA labeling of bacteria with proteins. SpyTag(*st*)-containing peptides are equipped with a SrtA-recognition motif (LPETGG) and mixed with bacteria and exogenous SrtA. In a second step, bacteria-labeled with *st* peptides are incubated with GFP-SpyCatcher (*sc*). (**f**) *S. epi* LM087 was incubated with or without *st*-LPETG, GFP-*sc* and SrtA. Conjugation of GFP to *S. epi* is measured by flow cytometry. (**g**) Representative microscopy images of SrtA labeling of *S. epi* LM087 with GFP-*sc* using SrtA. (**h**) Panel of Gram-positive bacteria derived from the microbiome labeled with 500μM of peptide (Cy5-LPETGG in blue or Cy5-GEGTLP in purple) and 50 μM SrtA. (c,d,f,g,h) Representative of three independent experiments.

We hypothesized that the natural process used by Gram-positive bacteria to anchor proteins to their cell-wall^6^ could be harnessed to display antigens on their surface^7,8^. Antigens—in the form of purified proteins or peptides—would be equipped with a short amino acid recognition site (LPETG^6^) for anchoring to the cell wall by the transpeptidase sortase A (SrtA) (Fig. 1b).

To test the feasibility of the approach, we first confirmed labeling of the *S. epi* cell wall using a fluorescent peptide, Cy5-LPETGG. As a negative control, bacteria were incubated with a scrambled version (Cy5-GEGTLP) not recognized by SrtA^9^. Because the native sortase expressed by *S. epi* was not sufficient to conjugate LPETG-containing peptides, we added exogeneous SrtA, a heptamutant version of *Staphylococcus aureus* SrtA with enhanced catalytic activity to boost the rate of the reaction (Extended Data Fig. 2a)^10^. This experiment confirmed that *S. epi* could be efficiently labeled with Cy5-LPETGG, but not Cy5-GEGTLP (Fig. 1c-d and Extended Data Fig. 2a-b). By varying the concentrations of peptide and enzyme, we determined that 500 μM peptide and 50 μM SrtA were optimal (Extended Data Fig. 2c), consistent with previous literature^11^.

Next, we asked whether a larger substrate—a folded protein—could also be attached to the cell surface in a similar fashion. Initial attempts to conjugate proteins with SrtA had negligible efficiency (Data not shown), presumably due to poor accessibility of large proteins to the Lipid II-based nucleophiles present in the bacterial cell wall. To circumvent these limitations, we adapted the SpyCatcher system^5,12^ by labeling the cell surface with peptides containing a SpyTag (*st*) and conjugating a SpyCatcher (*sc*) fusion protein to the *st* containing-peptide (Fig. 1e). This approach led to efficient attachment of GFP, and confirmed that all of the components of the reaction were required (Fig. 1f-g and Extended Data Fig. 2d). Finally, we examined the breadth of the SrtA labeling approach by treating a panel of Gram-positive bacteria from the murine and human microbiome with SrtA and a Cy5-containing peptide (Fig. 1h). Labeling was broadly effective, indicating that this method could facilitate the study of immune modulation by a range of genetically intractable colonists.

To illustrate the utility of this approach, we focused on *Cutibacterium acnes*, a highly prevalent skin commensal^13^ with limited genetic tools^14^. First, we asked whether unlabeled strains of *Cutibacterium* modulate the immune system and induce an antibody response upon colonization. Specific pathogen-free (SPF) mice were colonized topically and intranasally with a panel of *Cutibacterium*, including strains of *C. acnes* and *Cutibacterium avidum*. After 6 weeks, the serum IgG titers against each of the strains were high (Extended Data Fig. 3a-c). The response was cross-reactive among *C. acnes* strains but not with *C. avidum* (Extended Data Fig. 3d). This contrasts to the response to *S. epi*, for which cross reactivity was only observed between members of the same clade^2^. As with *S. epi*, the systemic antibody response to *Cutibacterium* was accompanied by a mucosal IgA response in the respiratory tract; interestingly, we also observed high IgA titers in the gastrointestinal and vaginal tract (Extended Data Fig. 3e).

To determine whether the immunogenicity of *Cutibacterium* colonization is conserved in humans, we measured antibody responses in the serum of 11 healthy human donors against a panel of independently sourced *Cutibacterium* strains. We noted high IgG titers against *C. acnes* (Extended Data Fig. 3f), greater than what we had observed against *S. epi*^2^. This suggests that the antibody response to *Cutibacterium* spp., especially *C. acnes*, is particularly strong in humans.

We next asked whether strains known to stimulate the immune system could be labeled with immunogens and activate antigen-specific T and B cells. We adoptively transferred ovalbumin (OVA)-specific CD4^+^ (OTII) or CD8^+^ T cells (OTI) into congenically-marked SPF mice that were colonized with *S. epi* or *C. acnes* enzymatically labeled with OTI-LPETG or OTII-LPETG. After one week, we observed T cell proliferation in the draining lymph nodes of mice colonized with *S. epi* or *C. acnes* labeled with their cognate antigens (Fig. 2a and Extended Data Fig. 4). Proliferation induced by colonization with labeled bacteria was as potent as with *S. epi* expressing OVA antigens genetically. Thus, SrtA labeling is an efficient strategy to display antigens on the surface of commensals and activate T cells in vivo.

**Figure 2:**
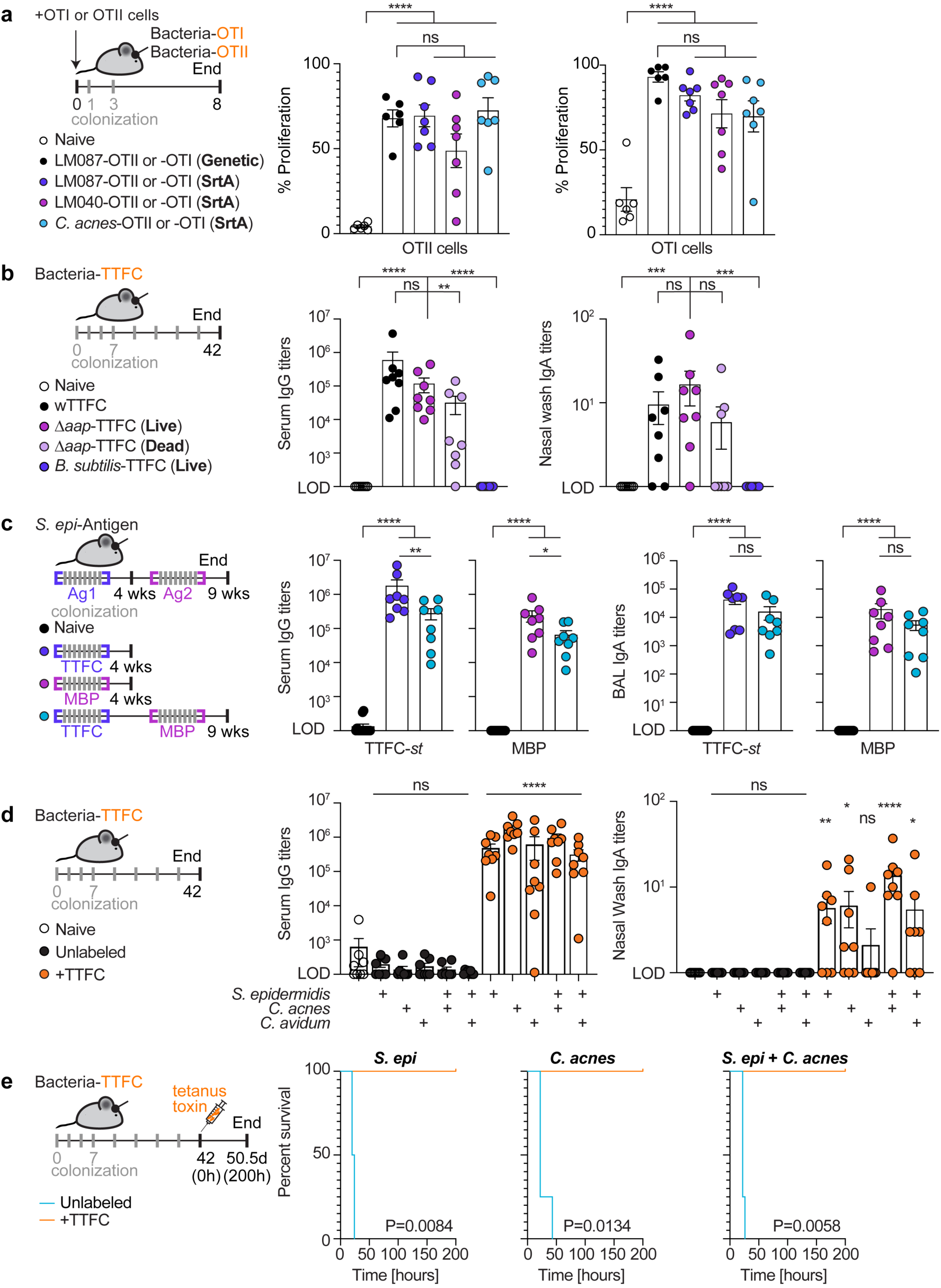
SrtA labeling can be used to induce T and B cell responses in vivo using bacteria with limited genetic tools. (**a**) 250,000 Cell-Trace Violet labeled Ovalbumin (OVA)-specific CD4^+^ (OTII) or CD8^+^ (OTI) T cells were transferred into congenically-marked SPF mice (day 0). On days 1 and 3, mice were colonized with bacteria displaying a OTII or OTI epitope on their wall by recombinant expression (“Genetic”) or by SrtA-mediated labeling (“SrtA”). LM087 is a genetically tractable strain of *S. epi* while LM040 is an intractable strain of *S. epi*^4^. Proliferation was measured by dilution of Cell-Trace Violet dye in the cervical draining lymph nodes by flow cytometry at day 8. (**b**) SPF mice were colonized with *S. epi* expressing a tetanus toxin fragment C (TTFC) fusion on its surface (wTTFC), live or heat-killed *S. epi* strain Δ*aap* labeled with TTFC using SrtA (*st*-LPETG + SrtA + TTFC-*sc*) or the soil bacterium *B. subtilis* labeled with TTFC. After 6 weeks, TTFC-specific antibody titers were measured in the serum and nasal washes by ELISA. (**c**) SPF mice were colonized with *S. epi* labeled with TTFC, maltose-binding protein (MBP) or TTFC and then MBP using the SrtA-mediated approach. Antibody titers to TTFC and MBP were measured at 4 or 9 weeks post-colonization in the serum and bronchoalveolar lavage (BAL) fluid. Glycerol stocks of labeled bacteria were used for each of the 8 colonizations per antigen. (**d**) SPF mice were colonized with *S. epi*, *C. acnes* or *C. avidum* or a combination of strains unlabeled or labeled with TTFC (SrtA approach). After 6 weeks, antibody titers against TTFC were measured in the serum and nasal washes by ELISA. (**e**) SPF mice were colonized with *S. epi*, *C. acnes* or a combination of both strains labeled or not with TTFC (SrtA approach). After 6 weeks, the mice were challenged with a lethal dose of tetanus toxin. n=4 except for the *C. acnes*-TTFC group (n=3). (a-d) Shows all mice from two independent experiments pooled and (**e**) all mice from a single experiment. P values were calculated using One-way ANOVA with Sidak’s multiple comparisons test (a), Dunnett’s (b and d) or Tukey’s (c) and log-rank tests (e). LOD=limit of detection.

Next, we asked whether the approach could also be used to induce antibody responses in vivo. SPF mice were colonized over 6 weeks with bacteria enzymatically labeled with tetanus toxin fragment C (TTFC), a non-toxic fragment of the neurotoxin tetanus toxin^15^ (Extended Data Fig. 5a). Colonization with *S. epi* labeled with TTFC led to strong antibody titers, as potent as a strain genetically engineered to express the same antigen (wTTFC). The response was observed systemically (serum) and across all mucosal sites sampled (nose, lungs, small and large intestine and vaginal tract) (Fig. 2b and Extended Data Fig. 5b). Taking advantage of the observation that antigens can be attached enzymatically to dead bacteria (Extended Data Fig. 5c-d), we used this approach to ask whether bacteria need to be alive to induce an antibody response in vivo. Mice associated with heat-killed *S. epi* labeled with TTFC had weaker and more variable antibody responses (Fig. 2b and Extended Data Fig. 5b), suggesting that bacterial viability contributes to the antibody response. Similarly, the soil bacterium *B. subtilis*—labeled enzymatically with TTFC—yielded no detectable response, despite efficient conjugation with SrtA (Fig. 2b and Extended Data Fig. 5b and e). Overall, these data show that the SrtA labeling approach can be used to induce robust B cell responses in vivo and characterize the requirements of this process.

The efficiency of homologous prime-boost regimens, where the same vector is reused multiple times with distinct antigens, often declines due to the immunity generated against the vector^16^. We wondered whether a skin commensal could be reused to induce immunity against a distinct antigen or if repeated use of the same bacterial strain would result in similarly waning titers. First, we ensured that responses could be elicited against antigens beyond TTFC. We used maltose binding protein (MBP) as a model antigen, taking advantage of its stability, immunogenicity and ease of expression^17^. Mice colonized with *S. epi* labeled with MBP had high titers of MBP-specific antibodies systemically and at mucosal sites (Fig. 2c and Extended Data Fig. 6). Encouraged by these results, we next tested whether *S. epi* could be reused for a second round of immunization. Mice were colonized with *S. epi* labeled with TTFC. After 4 weeks of colonization, as expected, we observed high titers of antibodies against TTFC (Fig. 2c and Extended Data Fig. 6). A week later, another cohort of mice was colonized with *S. epi* labeled with MBP. At 9 weeks post colonization, we observed high antibody titers against both TTFC and MBP in the serum and across mucosal sites. While there was a slight reduction in the serum response to the second antigen (i.e. MBP), there was no difference at mucosal sites (Fig. 2c and Extended Data Fig. 6). Thus, SrtA labeling of the same commensal can be used to sequentially vaccinate against multiple antigens.

Next, we asked whether we could induce antibody responses using two different colonists simultaneously. Because *S. epi* and *C. acnes* typically co-colonize the same hair follicle^18^, we asked whether applying both strains simultaneously—labeled with the same antigen—would lead to an enhanced antibody response. We also included *C. avidum* in this panel of strains because it generated such a strong antibody response in mice (see Extended Data Fig. 3c). Mice colonized with *S. epi*, *C. acnes* or a combination of *S. epi* with either *Cutibacterium* strain all had high serum titers against TTFC (Fig. 2d and Extended Data fig. 7a). In contrast, colonization with *C. avidum* labeled with TTFC led to a poor response, possibly because the labeling reaction was not as effective (Extended Data Fig. 7b). Finally, we assessed whether the titers induced using SrtA-labeled bacteria led to antibody responses potent enough to protect in a challenge experiment (Fig. 2e and Extended Data Fig. 7c). Mice were colonized with combinations of *S. epi*, *C. acnes* and *C. avidum* labeled with TTFC over 6 weeks and challenged with a lethal dose of tetanus toxin. Mice colonized with *S. epi*, *C. acnes*, *C. avidum* or *S. epi* + *C. acnes* were all fully protected against tetanus toxin. This suggests that the SrtA labeling method is compatible with a range of skin organisms and evoked an antibody response sufficiently strong to protect against lethal challenge with tetanus toxin.

Together with previous work demonstrating antigen-specific tolerance through SrtA-mediated antigen attachment to erythrocytes^19^, our work establishes the use of recombinant SrtA as a straightforward and powerful approach to display antigens on the surface of Gram-positive bacteria to modulate the immune system. The immunity generated by SrtA-labeled bacterial strains is comparable to genetically engineered strains and powerful enough to protect against a lethal dose of neurotoxin. By bypassing the need for genetic tools, our approach—and a complementary approach outlined in a companion study^20^—extend the repertoire of microbes that can be exploited to modulate host immunity. In the future, combinations of microbes could improve the magnitude, quality and durability of commensal vaccines and uncover how the microbiome naturally shapes immunity.

## METHODS

### Mice

Mice were housed under specific pathogen-free (SPF) conditions at Stanford University in a 12/12 light/dark cycle, at room temperature and a humidity level of 40-65% following to the guidelines established by the animal care and use committee. We purchased mice from Taconic (C57BL/6 (B6); SD, CC or Seattle facilities), OTI *rag2*^−/−^(#2334), OTII *rag2*^−/−^(#11490), and CD45.1 from Jackson (#002014). Mice were randomized at the beginning of each experiment. Experiments were not performed blindly, as the same person running the experiment was also performing the analysis.

### Bacterial cultures

*Staphylococcus epidermidis, Streptococcus spp., Staphylococcus xylosus and Bacillus subtilis* were grown on Difco Brain Heart Infusion (BHI) agar plates (BD 241810) overnight at 37 °C. Single colonies were then picked and inoculated into BHI broth (Difco Brain Heart Infusion media, BD 237200) and grown shaking overnight at 220 rpm. Strains carrying a pLI50-based plasmid were grown on/in BHI + chloramphenicol (10 µg/ml) (Sigma, C1919). *Cutibacterium* strains were grown anaerobically at 37 °C on pre-reduced 5% Columbia blood agar plates (VWR INTERNATIONAL, INC., A16). Single colonies were inoculated into pre-reduced BHI and grown for 3-4 days anaerobically at 37 °C. *Corynebacterium* strains were grown in BHI supplemented with 1% tween 80 shaking overnight at 220 rpm at 37 °C. *Lactobacillus* strains were grown in MRS standing at 37 °C.

### Bacterial colonization

As previously described^2^, 6-10-week-old C57BL/6 mice were colonized with skin commensals including *Staphylococcus epidermidis, Cutibacterium acnes, Cutibacterium avidum or B. subtilis* (See below for labeling procedures). Briefly, overnight cultures were OD normalized to an OD_600_ = 6 with BHI or BHI + chloramphenicol (10 µg/ml) where appropriate (Sigma) and gently applied to the ears, head and nostrils with sterile cotton swabs (Puritan, 25-8062PD) following the colonization schedule indicated in the figures.

### Sample collection from mice

At the end of the experiment, mice were sacrificed and samples harvested, as previously described^2^. To collect serum, the blood was harvested by cardiac puncture and the serum separated using serum tubes (BD, 365967) according to the manufacturer’s guidelines. The nasal washes were harvested by flushing 1 ml of PBS up to the trachea into the nasal cavity using 1 ml syringe attached to a 22G catheter (Exel International, 26746). The bronchoalveolar lavages (BAL) were collected by inserting a precut feeding tube (Instech Laboratories, FTP2038) down the trachea. After securing the feeding tube by tightening a knot with cotton string (Amazon, B08VWHH8LC) around the trachea, the lungs were flushed with 2x 1 ml of PBS. Small and large intestine contents were collected by emptying the organs using forceps on a paper towel prewetted with PBS and normalized to 100 mg/ml in PBS containing protease inhibitor (1 tablet in 50 ml of PBS, Sigma #11836170001). Intestinal contents were vortexed for 5 min in PBS + protease inhibitor solution, spun down at 10,000 g at 4 °C for 5 min and the supernatants transferred to fresh tubes and stored at −80 °C. Vaginal washes were harvested by flushing 20 μl of PBS 3x and pooling the samples obtained on 3 consecutive days.

### Adoptive transfer experiment

To isolate monoclonal T cells, spleens from the indicated mice were harvested, smashed on a 100 μm strainer (Miltenyi Biotec, 130-110-917) using the back of a 1 ml syringe (Becton Dickinson, 309659) in filter sterilized (MILLIPORE S2GPU05RE) PBE buffer (Miltenyi Biotec, one bottle of autoMACS Rinsing Solution supplemented with one bottle of BSA stock solution, 130-091-222 and 130-091-376). After spinning the splenocytes at 550 g for 5 min at 4 °C, the red blood cells were lysed using 1 ml of Ack lysis buffer for 5 min at room temperature (Thermo Scientific, # A1049201). T cells were then isolated using naïve CD4 and CD8 T cell isolation kits, where appropriate following manufacturing guidelines (Milteny, 130-104-453 or 130-096-543). T cells were then counted and rested for at least 30 min in complete RPMI (cRPMI = RPMI (Thermo Fisher, 61870127) supplemented with Penicillin-Streptomycin (Thermo Fisher, 15140122), HEPES (Thermo Fisher, 15630080), non-essential amino acids (Thermo Fisher, 11140050), sodium pyruvate (Thermo Fisher, 11360070), β-mercaptoethanol (Life Technologies Corporation, 21985023) and 10% FBS (GeminiBio)). For the Cell-Trace Violet labeling (Thermo, C34557), the cells were then washed twice with cold PBS (Thermo Fisher, 10010049) and resuspended in pre-warmed PBS at the concentration of 2 mio cells/ml and mixed 1:1 with a 2x staining mix of Cell-Trace Violet (10 μM for a 5 μM final concentration). The cells were labeled for 20 min in a 37 °C incubator protected from light. At the end of the incubation, cold cRPMI was added to the top of the 50 ml Falcon tube and spun down for 5 min at 4 °C at 550 g. The cells were washed one more time with cold cRPMI and resuspended into cRPMI and rested on ice for at least 30 min protected from light. During that time, the cells were counted. At the end of the resting period, the cells were spun down and washed 2 times with cold PBS and resuspended into cold PBS at a concentration of 2.5 mio cells/ml. Each mouse received 100 μl of cells (250,000 cells/mouse) via retroorbital intravenous injection. The next day, overnight cultures of *S. epi* and *C. acnes* (*C. acnes* was grown for 3.5 days at 37 °C in an anaerobic chamber) were OD normalized (1 cage = 5 ml of OD_600_=6) and washed twice with PBS at RT and resuspended into 500 μl of PBS and mixed 1:1 with 2x staining mix (1mM peptide + 100 μM SrtA for a final concentration of 500 μM peptide + 50μM SrtA in PBS). After 15 min of incubation at 4 °C rotating, the cells were spun down at 7,000 g at 4 °C for 3 min and washed 5 times with 1ml of PBS. The cells were resuspended into 5 ml of BHI and the mice colonized. The colonization was repeated on day 3. On day 8, the cervical draining lymph nodes were harvested, stained for surface markers and analyzed by flow cytometry (see below “Flow cytometry of mammalian cells”).

### Flow cytometry of mammalian cells

Draining cervical lymph nodes were isolated from mice and the cells extracted by smashing the lymph nodes on a 100 μm cell strainer (Miltenyi Biotec, 130-110-917) using the back of a 1 ml syringe (Becton Dickinson, 309659) in filter sterilized (MILLIPORE S2GPU05RE) PBE buffer (Miltenyi Biotec, one bottle of autoMACS Rinsing Solution supplemented with one bottle of BSA stock solution, 130-091-222 and 130-091-376). The dissociated cells were stained with antibodies raised against surface markers at 4 °C for 30 min in PBE, rinsed 2x with 200μl of PBE and fixed with BD cytofix/cytoperm according to manufacturer instructions. The cells were analyzed by flow cytometry on a FACSymphony machine (Becton Dickinson) and analyzed using FlowJo version 10.9.0 software package (Tree-Star).

### Sortase labeling with peptides for in vitro experiments

Overnight cultures of the indicated bacteria were harvested and the OD_600_ measured. An equivalent of 500 μl of OD_600_ = 6 per condition was transferred to a 1.5 ml Eppendorf, spun down at 14,000 g for 3 min at 4 °C and washed two times with PBS. The pellet was then resuspended in 100 μl of Sortase A (SrtA) labeling reaction containing 500 μM of peptide and 50 μM of SrtA in PBS and incubated for 15 min at 4 °C rotating. The bacteria were washed by adding 1ml of PBS and spun down at 14,000g for 3 min at 4 °C. The pellet was then resuspended into 100 μl of Syto9 staining buffer (1.5ul of Syto9 in 1ml of PBS, Thermo Fisher #S34854) and incubated for 15 min at 4 °C rotating. The bacteria were then washed two times with 1 ml of PBS, resuspended into 200 μl of PBS and analyzed by flow cytometry on a FACSymphony machine (Becton Dickinson) or a CytoFLEX machine (Beckman Coulter). The data was analyzed using FlowJo version 10.9.0 software package (Tree-Star).

### Sortase labeling with proteins for in vitro experiments

An equivalent of 500 μl of overnight cultures OD_600_ = 6 per condition was transferred to a 1.5 ml Eppendorf, spun down at 14,000 g for 3 min at 4 °C and washed two times with PBS. The pellet was then resuspended in 50 μl of PBS and 50 μl of 2x peptide staining mix (1mM peptide + 100μM in PBS for a final concentration of 500 μM of peptide and 50 μM of SrtA) was added and incubated for 15 min at 4 °C rotating. The pellet was washed 3x with 1 ml of PBS, resuspended into 50 μl of PBS and 50 μl of 2x protein staining mix was added (10mg/ml of GFP-*sc* in PBS) and incubated for 15 min at 4 °C rotating. The bacteria were washed with 1ml of PBS, resuspended into 100 μl of MycoLight™ Red JJ94 (AAT Bioquest) staining buffer (1:100 dilution) and incubated for 15 min at RT. The bacteria were washed two more times with 1 ml of PBS, resuspended into 200 μl of PBS and analyzed by flow cytometry on a FACSymphony machine (Becton Dickinson) or a CytoFLEX machine (Beckman Coulter). The data was analyzed using FlowJo version 10.9.0 software package (Tree-Star).

### Sortase labeling with proteins for in vivo experiments

An equivalent of 5 ml of OD_600_ = 6 per cage of overnight cultures of *S. epidermidis*, or *B. subtilis* was transferred into a 50 ml Falcon tube and diluted with up to 50 ml of PBS (fill to the top with PBS) and spun down at 3,600g for 8 min at RT. For “dead” *S. epi*, the bacteria were incubated for 30 min at 95 °C before proceeding to the next step. The pellet was then resuspended into 50 ml of PBS, spun down and resuspended into 500 μl of cold PBS and mixed with 500 μl of 2x staining solution containing 1mM of *st*-LPETG peptide and 100 μM of SrtA in PBS (final concentration: 500 μM of peptide and 50 μM of SrtA) and incubated for 15 min at 4 °C rotating. At the end of the incubation, the cells were spun down at 7,000g for 3 min at 4 °C, washed 5 times with 1 ml of cold PBS and resuspended into 500 μl of cold PBS and mixed 1:1 with 2x staining mix of TTFC-*sc* or MBP-*sc* (10 mg/ml for a final concentration of 5 mg/ml). After 15 min at 4 °C rotating, the cells were spun down at 7,000g for 3 min at 4 °C and washed 5x with 1 ml of cold PBS. The cells were then resuspended into 5ml of BHI and used for colonization. For *C. acnes* or *C. avidum*, the protocol was performed in the anaerobic chamber, and all the spins were performed at RT and the incubation on ice rather than at 4 °C for practical reasons. To prepare glycerol stocks, the procedure was the same except we prepared 10 cages at a time. The labeling reactions were performed in five 2ml Eppendorfs with 2 cages/tube. At the end of the labeling procedure, the bacteria were resuspended into 5 ml of BHI and combined into 1 falcon tube and mixed with 5 ml of 50% glycerol in BHI (final concentration 25% glycerol in BHI) and aliquoted (1 ml mix/tube) and stored at −80 °C. On the day of colonization, each tube was kept on dry ice and resuspended right before the colonization by adding 4 ml of RT BHI and used to colonize one cage.

### Bacterial ELISA

To measure antibody titers against whole bacteria, high-binding plates (Corning, 9018) were coated with 100 μl of poly-L-lysine solution (Sigma-Aldrich, P4707-50ML) for 15 min at room temperature, washed 3x with PBS, dried and coated with prewashed bacteria. To prepare the bacteria, overnight bacterial cultures were spun down at 3,600 g for 8 min at room temperature and washed two times with PBS (Thermo Fisher Scientific, 10010049) and normalized to an OD_600_ = 6 in PBS. ELISA plates were coated with 50 μl of normalized cultures per well, sealed (Bio-Rad, MSB1001) and incubated overnight at 4 °C on a flat surface. The following day, the plates were washed five times with PBST (PBS + 0.05% Tween-20, Sigma, P1379) before being blocked for 2h with 200 μl of 2% BSA (Sigma-Aldrich, A7030-100G) in PBST at room temperature. Next, the plates were incubated with sera from mice or healthy donors in serial dilutions. We typically used a first dilution of 1:100 (1:1000 for human sera against *Cutibacterium*) and loaded 100 μl (for 1:3 dilutions) or 90 μl (for 1:4 dilutions) and serially diluted the samples (using 2%BSA/PBST as dilution buffer). For nasal washes, BAL IgA, and intestinal contents we loaded undiluted samples and for the vaginal washes we used an initial 1:3 dilution, as indicated in the figure (i.e see limit of detection = LOD). After 1 h of incubation at RT, the plates were washed five times with PBST and incubated with secondary antibody for 1 h at a dilution of 1:10,000 (mouse IgG: SouthernBiotech, 1030-05; Mouse IgA: SouthernBiotech, 1040-05, Human IgG: Thermo Fisher, 31413) in 2% BSA/PBST. The plates were then washed 5 more times with PBST, revealed for 30 min with 100 μl of TMB reagents (BD, 555214) and the reaction stopped with 50 μl of stop solution (BioLegend, 423001). Antibody titers were calculated by logarithmic interpolation of the dilutions with absorbance at 450 nm immediately above and below the absolute value 0.2 (after subtracting the background), as previously described^21^.

### Cloning

For all constructs, the backbones and inserts were amplified using PrimeStar MAX DNA Polymerase (Takara Bio, R045B) and purified using a PCR cleanup kit (Zymo, D4014). If a plasmid was used as template for the backbone, we digested the PCR product with Dpn1 (NEB, R0176L) to eliminate the template DNA, following manufacturer’s guidelines. Purified inserts and backbones were then ligated using Gibson assembly (NEB, E2621L) and transformed into stable competent cells (NEB, C3040H) for pET expression vectors or chemically competent DC10B for pLI50 vectors, similarly to^2,4^. Briefly, for pLI50 vectors, 3 μl of a 10 μl Gibson reaction was added to chemically competent DC10B cells, incubated for 30 min on ice, heat-shocked at 42°C for 45 sec and recovered for 1h in LB at 37 °C shaking. The cells were then plated onto LB + Carbenicillin 50=50 μg/ml (Teknova, C2105). After sequence confirmation, the vectors used for bacterial expression (pET vectors) were transformed into competent BL21 (DE3) (NEB, C2527H) or *S. epi* strain LM087 for pLI50 vectors, as described previously^2^.

### Protein production and purification in bacteria

Superfolder-GFP (sGFP)-SpyCatcher003, and TTFC-SpyCatcher003 were cloned into a pET-30b expression vector while TTFC-SpyTag003 and MBP-SpyCatcher003 into a pET-30b expression vector (see above) and transformed into BL21 (DE3) strain (NEB, C2527H) for expression and protein purification as described before^2^. SrtA expression plasmid was obtained from Addgene (#51141). Briefly, single colonies were inoculated into LB containing kanamycin (50 μg/ml) for overnight growth at 37 °C with shaking at 220 rpm. The next day, the cultures were back diluted ∼1:200 times into 1 L of Terrific Broth (Fisher Scientific, AAJ75856A1) + Kanamycin (50 μg/ml). When the cultures reached OD_600_ = ∼ 0.4 - 0.6, protein expression was induced by adding 1 mM IPTG and the cultures transferred to 28 °C for overnight induction. The next day, the bacteria were harvested by spinning the cultures at 3,600 x *g* for 20 min at 4 °C and resuspended into 30 ml of ice-cold lysis buffer (50mM Tris-HCl pH 7.5 + 150 mM NaCl + 10 mM imidazole) containing DNAse (Sigma, 10104159001), lysozyme (Sigma, L6876) and protease inhibitor (Sigma, 11836170001, 1 tablet for every 50 ml of buffer). The pellet was then sonicated 2-3x for 5 min (15 sec sonication at 40 % output, stop 45 sec, Fischer Scientific model FB120, 0.63cm probe), cleared by centrifuging for 1.5 h at 20,000 x *g* at 4 °C. The clarified lysate was then incubated for 1 h with 6 ml of prewashed Ni-NTA beads rotating at 4 °C. The beads were loaded onto polypropylene columns (Qiagen, 34964, 2 columns/L of starting culture), washed four times with 25 ml of lysis buffer per column and eluted in 8 ml of elution buffer (4 ml per column) (50 mM Tris-HCl pH 7.5, 150 mM NaCl, 500 mM Imidazole). The eluate was concentrated on Amicon columns (EMD Millipore, UFC901024 or UFC903024 where appropriate), buffer exchanged on PD-10 columns (Cytiva, 17-0851-01, according to manufacturer’s instructions) into 50 mM Tris-HCl pH 7.5 + 150mM NaCl containing 10 % of glycerol, aliquoted, flash frozen and stored at −80 °C. Protein purity was assessed by Coomassie staining (Fisher Scientific, SimplyBlue™ SafeStain, LC6060).

### Protein production and purification in mammalian cells

MBP expression used for ELISA was carried out in Expi293 suspension cells maintained at 37 °C under 8% CO_2_ and >80% humidity, shaking at 125–130 rpm on a 19 mm orbital shaker. On the day of transfection, log-phase cultures were adjusted to 2.5 × 10⁶ viable cells/mL in non-baffled flasks. Transfection complexes were formed by diluting transfection-grade plasmid DNA (1 μg per mL of culture) in Opti-MEM (Thermo Fisher Scientific 31985088) with a three-minute room-temperature incubation, and separately diluting ExpiFectamine 293 Transfection Reagent (Thermo Fisher A14525, 2.7 μL per mL of culture) in Opti-MEM. The two solutions were combined and, after a five-minute incubation, added directly to the cell suspension; ExpiFectamine 293 Transfection Enhancers 1 and 2 were introduced 16–18 hours later. Four days post-transfection, culture supernatant containing the secreted, His-tagged MBP was collected by centrifugation (3,500 g, 15 min, 4 °C). The clarified supernatant was treated with PMSF (0.2M stock at 100x) and sodium azide (20% stock at 1000x) and passed through a 0.22 µm filter prior to purification. Protein was purified by batch affinity chromatography using fresh, washed nickel resin (Qiagen 30210), with gentle agitation for 1 h at 4 °C. The resin was washed and protein eluted with a 250 mM imidazole elution buffer. Eluate was buffer-exchanged into the desired final buffer using PD-10 desalting columns (Cytiva, 17-0851-01), concentrating in Amicon spin filters beforehand if needed (EMD Millipore, UFC901024). Final protein concentration was determined by nanodrop measurement or Qubit fluorometric assay, and purity was confirmed by Coomassie-staining of SDS-PAGE. Final aliquots were flash frozen and stored at −80 °C until use.

### Confocal microscopy of bacteria

*S. epidermidis* was grown on BHI agar plates overnight and BHI liquid culture overnight at 37 °C shaking under aerobic conditions. *C. acnes* and *C. avidum* were grown on Columbia agar plates supplemented with sheep blood for 4 days and in BHI liquid culture for 3 days at 37 °C in an anaerobic chamber. On the day of the experiment, 500 μL of OD_600_ = 6 normalized culture was pelleted and washed thrice with 1 mL PBS. For attachment of Cy5-conjugated peptides to bacteria, 100 μL reactions were performed with 50 μM SrtA and 200 µM Cy5-conjugated peptide for 20 minutes at room temperature. For attachment of GFP to bacteria, cells were first reacted with 200 μM SpyTag-LPETGG peptide and 50 μM SrtA for 20 minutes at room temperature in a volume of 100 μL. Then, cells were washed three times with 1 mL PBS and incubated for 30 minutes with 200 μL of 5 mg/mL SpyCatcher-GFP. All cells were subsequently washed three times with 1 mL PBS, before being stained in 100 μL at room temperature for 10 minutes with either MycoLight Red (AAT Bioquest 24006) or Syto9 (Invitrogen S34854), each used at a 1:100 final dilution. After two more washes, cells were fixed in 1 mL of 4% paraformaldehyde diluted in PBS for 10 minutes at room temperature while rocking. Cell pellets were again washed three times in 1 mL PBS before finally being resuspended in 800 μL PBS. Cover slips were coated with 0.1% poly-lysine (Sigma-Aldrich P8920-500ML) at room temperature for 15 minutes and were then washed twice with water and dried. Bacterial cells were then added to cover slips and allowed to adhere for 10 minutes, washed with PBS, mounted using ProLong Gold Antifade Mountant (Thermo Scientific P36930), and sealed with nail polish. Slides were incubated overnight at 4 °C to cure. Images were acquired using a Zeiss LSM 880 confocal microscope using an AiryScan detector.

### Protein ELISA

ELISA high-binding plates (Corning, 9018) were coated overnight at 4 °C with 0.25 µg of TTFC-SpyTag003 or MBP (from mammalian cell production) per well in PBS on a flat surface. On the next day, the plates were washed 5x with PBST (PBS + 0.05% Tween-20, Sigma, P1379), blocked with 200 μl of 2% BSA (Sigma-Aldrich, A7030-100G) in PBST at room temperature for 2 h. The samples were then loaded and incubated for 1 h at RT. We typically loaded either 100 μl and then did 1:3 dilutions or 90 μl and performed 1:4 dilutions. The plates were then washed 5x with PBST and incubated with secondary antibody at a dilution of 1:10,000 (SouthernBiotech, 1030-05 or 1040-05) in 2 % BSA/PBST. After another 1 h incubation, the plates were washed 5 more times with PBST and developed using 100 μl of TMB reagents (BD, 555214) and the reaction stopped with 50 μl of stop solution after 30 min (BioLegend, 423001). Antibody titers were calculated by logarithmic interpolation of the dilutions with absorbance at 450 nm immediately above and below the absolute value 0.2, as previously described^21^.

### Tetanus toxin challenge experiment

The tetanus challenge experiments were performed following previously published protocols^2^. Briefly, tetanus toxin (List Biological Laboratories, 190A) was reconstituted in 250 μl of sterile water according to manufacturer’s instructions and diluted in dilution buffer = 0.9% NaCl (Fisher Scientific, ALX-505-009-LD15) plus 1 mg/ml mouse serum albumin (Sigma, A3139) at a concentration of 50 μg/ml, aliquoted and stored at −80 °C. On the experimental day, a frozen aliquot of toxin was thawed to room temperature for 1.5 h and diluted in dilution buffer to 800 pg/μl (4000 ng/kg dose). The mice were weighed and injected with 4000ng/kg of tetanus toxin intraperitoneally (100 μl for a 20 g mouse). Following the injection, the mice were monitored for 200 h (the experimental endpoint). Mice were euthanized when they displayed moderate tetanus symptoms (hunched back and disappearance of the righting reflex) and considered positive for tetanus.

### Phylogenetic tree generation

The phylogenetic analysis of *Cutibacterium* strains was generated from full length 16S rRNA gene sequences using the nf phylogenetic tree workflow (https://github.com/FischbachLab/nf-phylogenetic-tree/). The pipeline includes three major steps: (1) multiple sequence alignment with MAFFT using the parameters --localpair --adjustdirectionaccurately --maxiterate 1000 --auto; (2) phylogenetic tree construction with RAxML (-m GTRGAMMA -p 12345 -f a -x 12345 -N 100); and (3) visualization and export of the final tree as a PDF using FigTree.

### Statistical analyses

All statistical analyses were performed using the Prism 11.0.1 software (GraphPad Software, La Jolla, CA). Unpaired two-tailed Student’s t tests were used when two groups were compared. When more than two groups were compared, a one-way ANOVA analysis was performed with appropriate post-hoc tests: Tukey’s when comparing all group combinations, Dunnett’s when one group is compared to all other groups or Šídák’s when specific comparisons were performed. When data was plotted on a log10 scale, a lognormal distribution was assumed. For survival experiments (tetanus challenge), a log-rank test was used. All tests used are indicated in the figure legends and the P values are reported in Supplemental Table S2. P values below 0.05 were considered statistically significant.

## EXTENDED DATA FIGURES

**Extended Data Figure 1:**
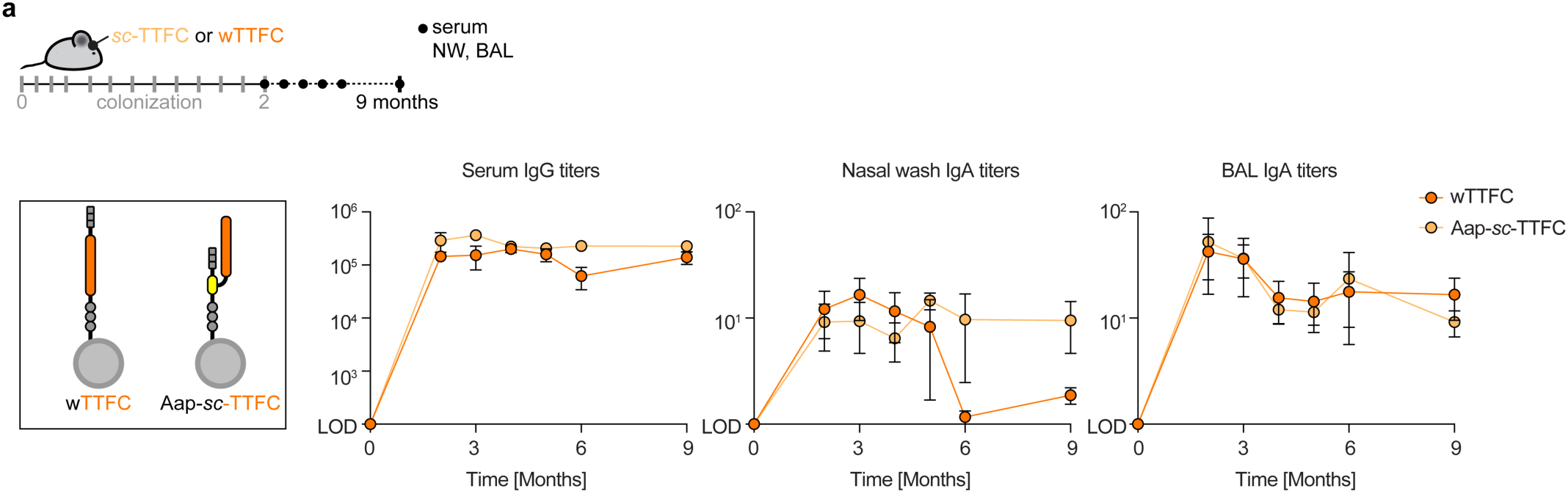
Colonization with engineered *S. epi* strains leads to long-lived antibody responses. (**a**) SPF mice were colonized over 6 weeks with *S. epi* displaying the model antigen TTFC on its surface by either genetic fusion (wTTFC) or by bioconjugation to a SpyCatcher (*sc*) motif fused to the wall protein Aap (Aap-*sc*-TTFC). Antibody titers were measured by ELISA. BAL=bronchoalveolar lavage. (n=3) mice per timepoint of one experiment. LOD=limit of detection.

**Extended Data Figure 2:**
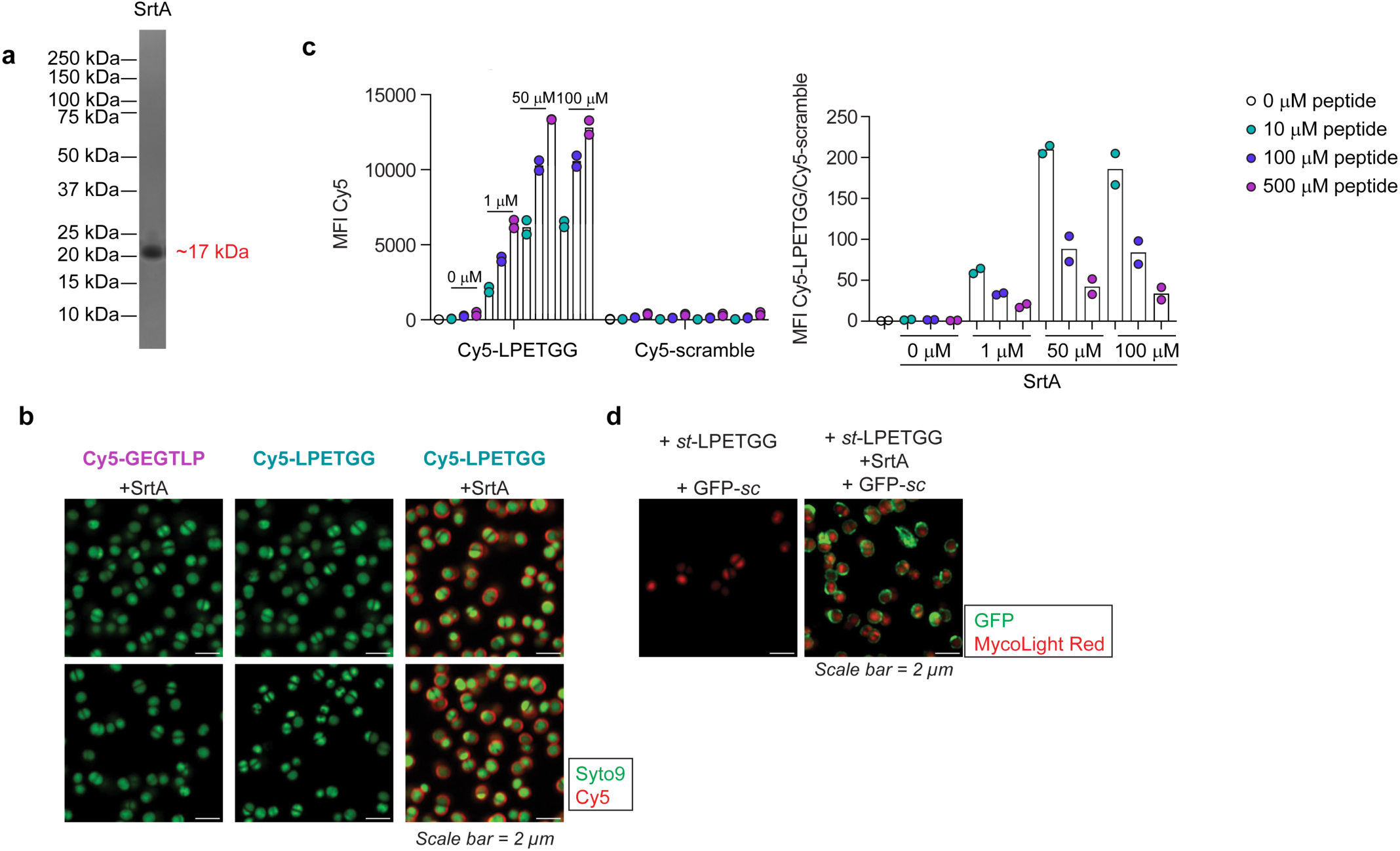
Optimization of the SrtA labeling method. (**a**) SrtA was purified from *E. coli* and the purity assessed by Coomassie. (**b**) Representative images of *S. epi* strain LM087 labeled with 500 μM of the indicated Cy5 peptides in the presence or absence of 50 μM of SrtA. (**c**) *S. epi* strain LM087 was labeled with different amounts of SrtA and Cy5 peptides containing either a SrtA recognition motif (LPETGG) or a scramble version of the same peptide (GEGTLP). Graphs show the labeling efficiency as measured by flow cytometry of two independent experiments, representative of three experiments. (**d**) Representative images of *S. epi* strain LM087 labeled with GFP-*sc* and *st*-LPETG in the presence or absence of 50 μM of SrtA. (b,d) Representative of three independent experiments.

**Extended Data Figure 3:**
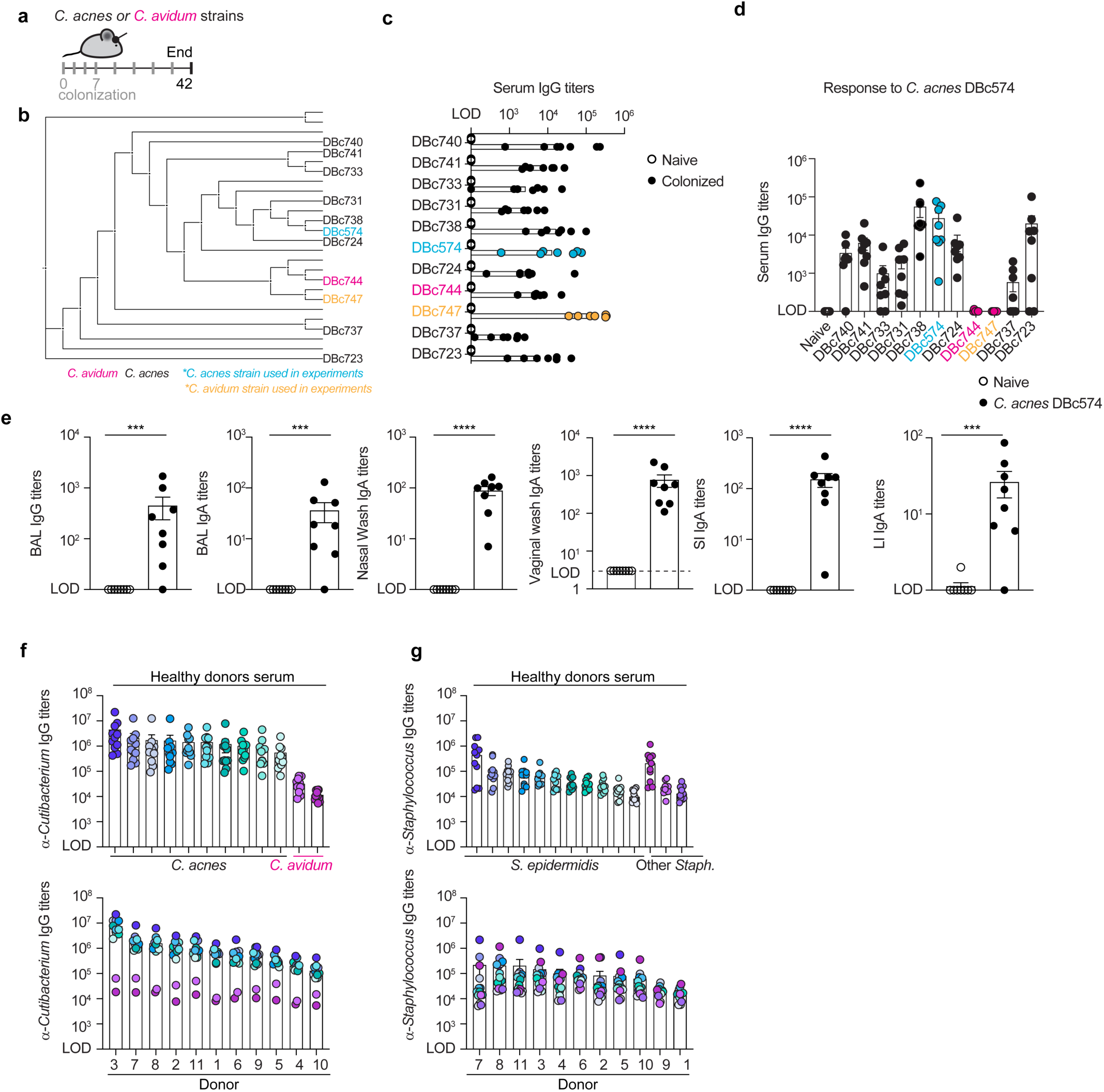
*Cutibacterium acnes* and *Cutibacterium avidum* induce strong antibody responses in mice and humans. (**a)** Schematic of the experiment for data shown in (c-e). SPF mice were colonized over 6 weeks with one strain of a panel of *C. acnes* or *C. avidum*. (**b**) Phylogenetic tree of a set of *Cutibacterium* strains based on 16s rRNA sequences. (**c**) The serum IgG titers against each bacterial strain were measured by ELISA for the experiment described in (a). (**d**) For the experiment shown in (a), the serum of mice colonized with different *Cutibacterium* strains was tested for reactivity against *C. acnes* strain DBc574, a primary isolate from a healthy donor. (**e**) Antibody titers across mucosal sites for mice colonized with *C. acnes* strain DBc574 or not colonized. BAL=bronchoalveolar lavages, SI=small intestine, LI=large intestine. P values were calculated using unpaired two-sided lognormal Student t tests. (**f**) IgG titers against a panel of *Cutibacterium* strains in the serum of eleven healthy donors. The data is either plotted by strain (top) or by donor (bottom). (**g**) For data adapted from ^2^, same as (f) but against a panel of *Staphylococcus* strains. (a-e) All mice shown from two independent experiments. LOD=limit of detection.

**Extended Data Figure 4:**
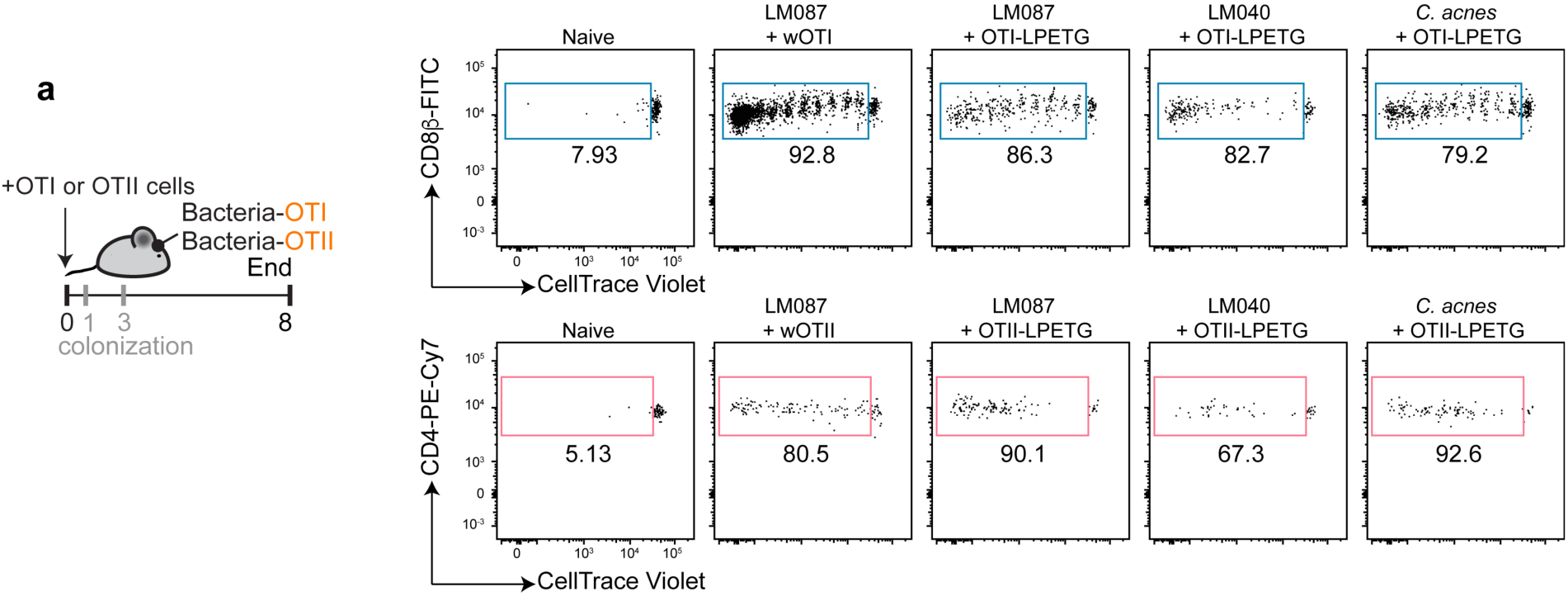
SrtA-labeled bacteria can prime CD4^+^ and CD8^+^ T cells in vivo. (**a**) For the experiment shown in Fig. 2a, representative flow plots from two independent experiments.

**Extended Data Figure 5:**
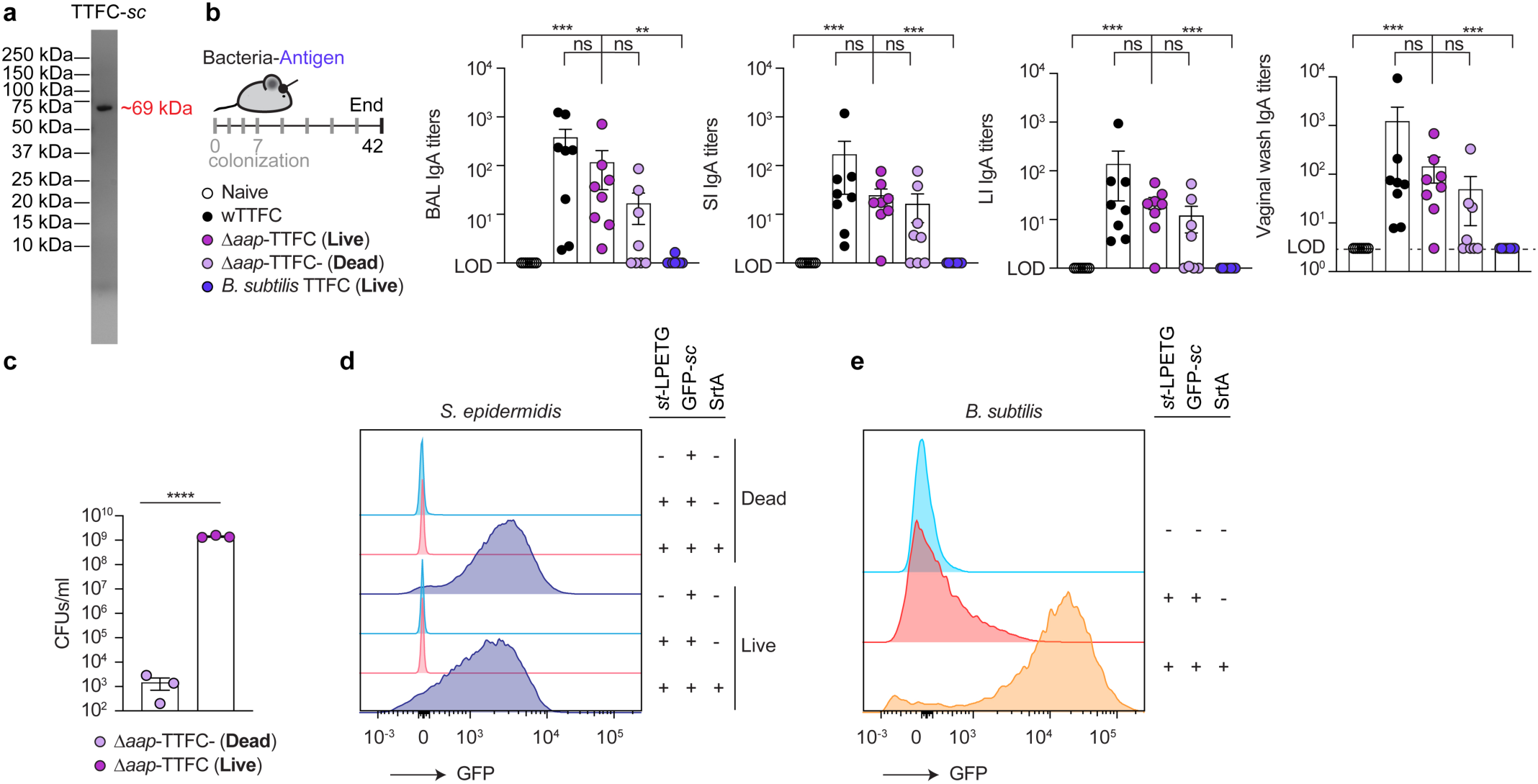
*S. epi* labeled with TTFC can induce a potent B cell response in vivo. (**a**) Representative Coomassie gel to assess the purity of TTFC-*sc* used in the in vivo experiments. (**b**) For the experiment shown in Fig. 2b, antibody titers against TTFC across mucosal sites. BAL=bronchoalveolar lavages, SI=small intestine, LI=large intestine. (**c**) Colony-forming Units (CFUs) for heat-killed *S. epi* Δ*aap* used in the above experiment. (**d-e**) Labeling efficiency of live and dead *S. epi* strain LM087 (d) or *B. subtilis* (e) as measured by flow cytometry. (b) shows all mice of two independent experiments, (c) three independent experiments shown and (d-e) representative of three independent experiments. P values were calculated using One-way ANOVA with Dunnett’s multiple comparisons tests (b) and an unpaired two-tailed lognormal Student’s t test (c). LOD=limit of detection.

**Extended Data Figure 6:**
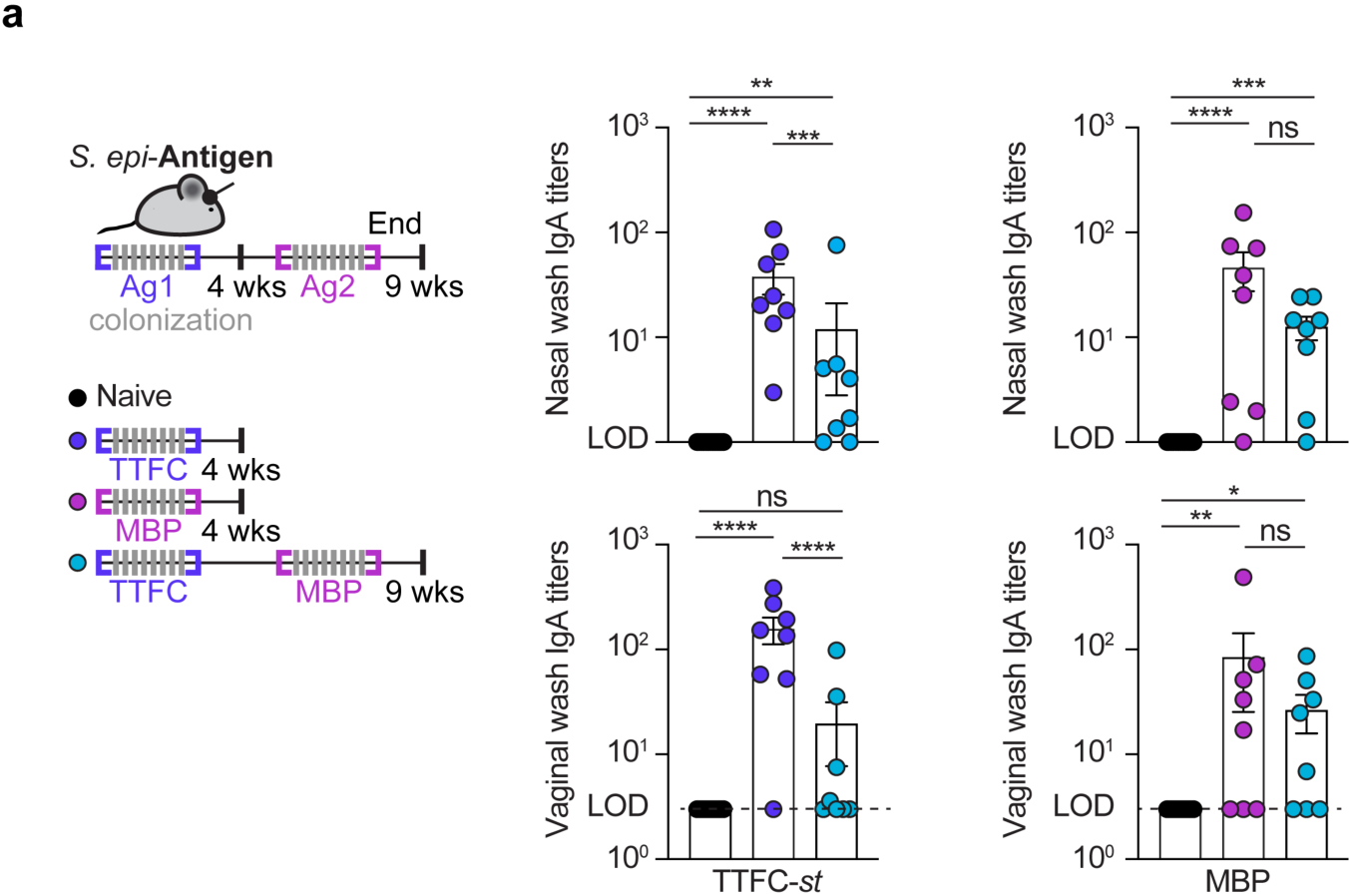
Induction of immunity against multiple antigens using SrtA labeling. (**a**) For experiment shown in Fig. 2c, antibody titers across mucosal sites as measured by ELISA against TTFC-*st* or MBP. All mice shown of two independent experiments. P values were calculated using One-way ANOVA with Tukey’s multiple comparisons tests. LOD=limit of detection.

**Extended Data Figure 7:**
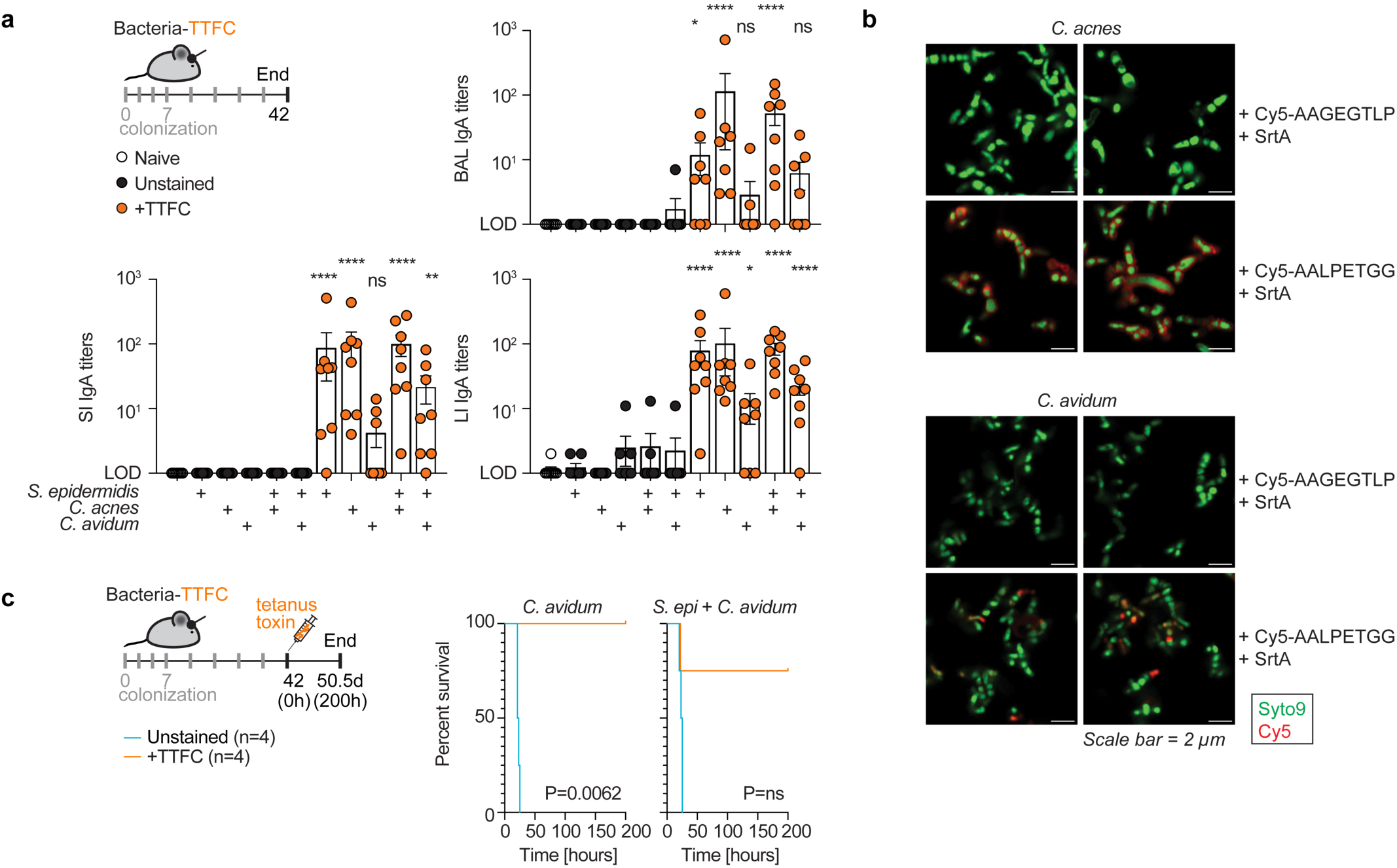
Colonization with antigen-labeled skin microbes leads to protective immunity in vivo. (**a**) For experiment shown in Fig. 2d, IgA titers in the bronchoalveolar lavages (BAL), small and large intestine (SI and LI). (**b**) Representative microscopy images of *C. acnes* and *C. avidum* labeled with Cy5-containing peptides with SrtA. (**c**) For experiment shown in Fig. 2e, survival curves after challenge with a lethal dose of tetanus toxin in mice colonized with SrtA-labeled *C. avidum* or *S. epi* + *C. avidum* versus unlabeled bacteria. (a) All mice shown from two independent experiments, (b) representative images of one experiment and (c) all mice shown of one experiment. P values were calculated using One-way ANOVA with Dunnett’s multiple comparisons tests (a) and log-rank tests (c). LOD=limit of detection.

## ACKNOWLEDGMENTS

We are deeply indebted to members of the Fischbach lab for helpful discussions and suggestions, especially Martin McLaughlin for chemical biology expertise, Amina Jbara for input on animal experiments and Xiandong Meng for assistance with phylogenetic analysis; Maria Prado, Amanda Espinoza and Joanne Au for keeping the lab running; Hidde Ploegh for careful review of the manuscript; Katherine Lemon and Barbara Bensing for sharing strains; members of the Stanford University Veterinary Service Center for animal husbandry, in particular Fa’atoto Veloz; and staff members of the Stanford University shared FACS facility for assistance with flow cytometry analysis (NIH grant 1S10OD026831-01). This work was supported by the Bill and Melinda Gates Foundation (M.A.F.), Coefficient Giving (M.A.F.), the Stanford Microbiome Therapies Initiative, the Swiss National Science Foundation (D.B., Early Postdoc.Mobility and Postdoc.Mobility), NIH grants 5R01AI175642-02 (M.A.F.), 1K99AI180358-01A1 (D.B.), the Leona M. and Harry B. Helmsley Charitable Trust (M.A.F.), the Chan Zuckerberg Biohub (M.A.F.), the Arc Institute (M.A.F.), Flu lab (M.A.F.), the Department of Defense NDSEG Fellowship (P.V.L.), the Knight-Hennessy Fellowship (P.V.L.), and a fellowship from The Helen Hay Whitney Foundation (K.D.B.).

## AUTHOR CONTRIBUTIONS

Conceptualization: D.B., and M.A.F., Methodology: D.B., K.D.B., P.V.L. and M.A.F., Investigation: D.B., K.D.B., P.V.L., E.T., T.T.D.N., M.C., A.J., A.Z., J.M.S., M.T., A.T., Visualization: D.B. and M.A.F., Funding acquisition: D.B., K.D.B., P.V.L., and M.A.F., Supervision: D.B. and M.A.F., Writing – original draft: D.B., Writing – review editing: D.B., K.D.B., P.V.L., E.T., T.T.D.N., A.J and M.A.F.

